# *In vivo* low-intensity magnetic pulses durably alter neocortical neuron excitability and spontaneous activity

**DOI:** 10.1101/2022.03.18.484911

**Authors:** Manon Boyer, Paul Baudin, Chloé Stengel, Antoni Valero-Cabre, Ann M. Lohof, Stéphane Charpier, Rachel M. Sherrard, Séverine Mahon

## Abstract

Magnetic brain stimulation is a promising treatment for neurological and psychiatric disorders. However, a better understanding of its effects at the individual neuron level is essential to improve its clinical application. We combined focal low-intensity repetitive transcranial magnetic stimulation (LI-rTMS) to the rat somatosensory cortex with intracellular recordings of subjacent pyramidal neurons *in vivo*. Continuous 10 Hz LI-rTMS reliably evoked firing at ∼4-5 Hz during the stimulation period and induced durable attenuation of synaptic activity and spontaneous firing in cortical neurons, through membrane hyperpolarization and a reduced intrinsic excitability. However, inducing firing in individual neurons by repeated intracellular current injection did not reproduce LI-rTMS effects on neuronal properties. These data provide novel understanding of mechanisms underlying magnetic brain stimulation showing that, in addition to inducing biochemical plasticity, even weak magnetic fields can activate neurons and enduringly modulate their excitability.

## Introduction

Repetitive transcranial magnetic stimulation (rTMS) is a promising tool for the treatment of neurological and psychiatric diseases. However, a better understanding of the cellular mechanisms underlying rTMS effects is needed to optimize current treatment protocols, whose outcomes remain variable (Bhattacharya et al., 2021; Cirillo et al., 2017; Rossini et al., 2015). Clinical application of magnetic stimulation delivers repeated high-intensity (0.5–2 Tesla, T) magnetic pulses to a targeted cortical area to induce an electric field that activates cortical neurons and initiates activity-dependent plasticity (Bhattacharya et al., 2021; Cirillo et al., 2017). Such plasticity is confirmed in rodents, where high-intensity magnetic stimulation *in vitro* induces synaptic plasticity in hippocampal networks (Lenz and Vlachos, 2016; Vlachos et al., 2012) and modulates neocortical inhibitory interneurons (Hoppenrath et al., 2016; Mix et al., 2015).

Besides high-field magnetic stimulation, whose duration and frequency of application is limited by technical and safety reasons (Rossini et al., 2015), new protocols using low-intensity (milliTesla, mT) magnetic fields are emerging. Human studies show that low-intensity stimulation to the entire brain can modulate EEG activity and cerebral function (Cook et al., 2009; Di Lazzaro et al., 2013; Rohan et al., 2014). In addition, focal application of these weak stimuli (low-intensity rTMS; LI-rTMS) in rodents facilitates the reorganisation of abnormal circuits (Makowiecki et al., 2014; Rodger et al., 2012), stimulates axonal growth and synaptogenesis (Dufor et al., 2019), and modifies cellular calcium signalling and gene expression (Dufor et al., 2019; Grehl et al., 2015). However, low-intensity stimulation is thought to generate electric fields weaker than those considered able to initiate action potential (AP) firing (Chan and Nicholson, 1986; Grehl et al., 2016; Radman et al., 2009). Thus, it remains unclear how LI-rTMS alters the properties of single neurons to contribute to structural and functional changes. Moreover, human cortical responses to magnetic stimulation critically depend on the ongoing activity of the stimulated brain region (Silvanto and Pascual-Leone, 2008). It is therefore necessary to identify the impact of LI-rTMS on the electrical properties of cortical neurons *in vivo, i*.*e*. in a physiologically relevant experimental condition that preserves neuronal connectivity.

We developed an *in vivo* model to deliver focal low-intensity (10 mT) rTMS to the somatosensory cortex (S1) of sedated rodents during intracellular recording of underlying pyramidal neurons. We provide direct evidence that LI-rTMS reliably elicits action potentials (APs) in S1 pyramidal cells. Also, 10 minutes of continuous 10 Hz LI-rTMS produces long lasting (≥ 40 minutes post-stimulation) modifications in intrinsic membrane properties and spontaneous activity. These after-effects include dampening of resting membrane properties, membrane hyperpolarization, and weaker background synaptic and firing activity. This was accompanied by altered transfer function, characterized by increased minimal current required to induce neuronal firing. These LI-rTMS-mediated effects were not reproduced by 10 minutes of AP firing, induced by intracellular injection of threshold current pulses at 10 Hz. Our data show that, like high-intensity stimulation, LI-rTMS activates neurons and durably modulates their activity and excitability. This changes the paradigm for brain stimulation with weak magnetic fields and increases the range of potentially therapeutic stimulation parameters for clinical use.

## Results

To study the immediate and time-dependent effects of LI-rTMS on cortical activity, we paired *in vivo* intracellular recordings of S1 neurons (*n*=41 from 31 rats) with ECoG monitoring (Fig. 1, A–C). Some recorded neurons were filled with neurobiotin (*n*=19) and confirmed histologically as pyramidal neurons located in layer 5 (*n*=11) or 2/3 (*n*=8; Fig. 1B). All recorded neurons displayed the characteristic firing of S1 pyramidal cells (Mahon and Charpier, 2012; Zhu and Connors, 1999): regular spiking (*n*=27; Fig. 3B) or intrinsic bursts of APs (*n*=14) following suprathreshold current pulse injection (Fig. 3B, fig. S1). Since no differences in LI-rTMS effects were observed between pyramidal neurons located in layers 2/3 or layer 5, we pooled the corresponding results.

**Fig. 1.**
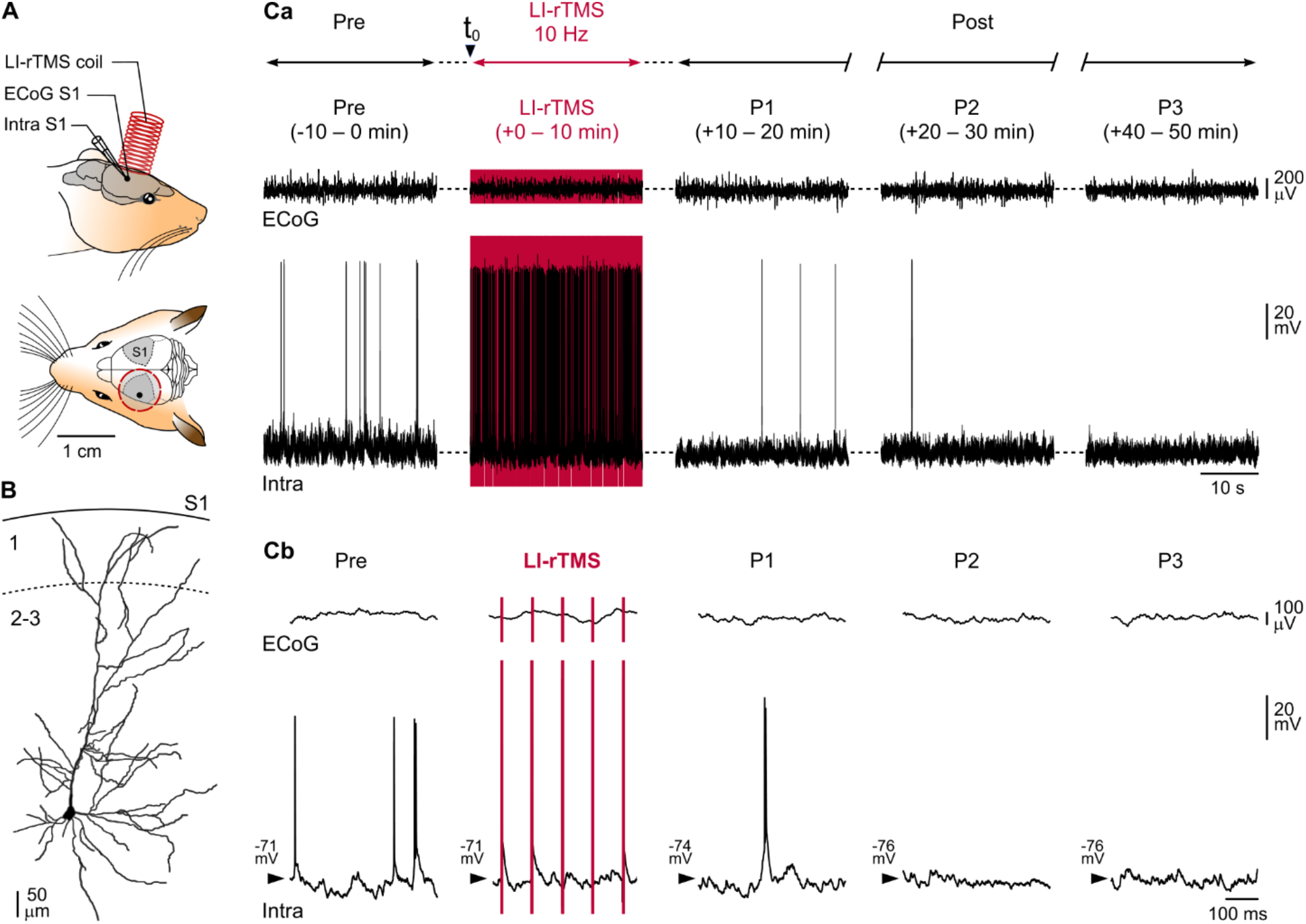
Experimental paradigm and definition of recording periods. (**A**) The experimental set-up illustrating the positions of the LI-rTMS coil and the ECoG (ECoG S1) and intracellular (Intra S1) recording electrodes. LI-rTMS (10 Hz for 10 minutes) was delivered through an 8-mm coil placed ∼2 mm above the brain surface. **(B)** Reconstruction of a layer 2/3 neurobiotin-filled pyramidal neuron **(Ca)** Simultaneous ECoG and intracellular recordings from a layer 2/3 pyramidal neuron, before (Pre), during LI-rTMS, and at different times (+10−20 min, P1; +20−30 min, P2; +40−50 min, P3) after the onset of LI-rTMS (Post). **(Cb)** Expanded view (500 ms) of the recordings shown in **(Ca)**. The LI-rTMS artifacts are shown in red. Here and in the following figures, the Vm is indicated at the left of intracellular recordings.

### LI-rTMS decreases spontaneous neuronal activity

First, we verified the stability of S1 cortical neuron recordings by measuring the variability of electrical membrane properties for 20 minutes preceding LI-rTMS (Fig. 1C, fig. S1). During this control period, the LI-rTMS coil was positioned above S1, but no stimulation was delivered. There were no significant changes in neuronal spontaneous activity or membrane excitability during this baseline period (fig. S1; see Supplementary Information for details). To analyze the time-dependent effects of LI-rTMS (*n*=16 experiments), we divided recordings into five consecutive periods: 1) pre-stimulation (Pre), the 10 minutes immediately before LI-rTMS; 2) 10 minutes LI-rTMS at 10 Hz; and 3) three successive post-stimulation times: 10–20 min (P1), 20–30 min (P2) and 40–50 min (P3) after the start of LI-rTMS (Fig. 1C). Before stimulation, spontaneous intracellular activity showed broadband frequency, small amplitude synaptic fluctuations, and a Gaussian-like distribution of membrane potential values (Vm) around -72.1 ± 4.8 mV (mean±SD; *n*=16 neurons) (Fig. 1C Pre, Fig. 2A-B). In half of the recorded neurons, background synaptic activity caused spontaneous firing at 1.8 ± 2.3 Hz (*n*=8) with high temporal variability (CV2 ISIs = 1.2 ± 0.3, *n*=8) (Figs. 1C, 2A). Consistent with this intracellular activity, the ECoG showed desynchronized, relatively fast, small-amplitude cortical waves (Fig. 1C), as classically observed under sufentanil sedation (Altwegg-Boussac et al., 2017, 2014).

**Fig. 2.**
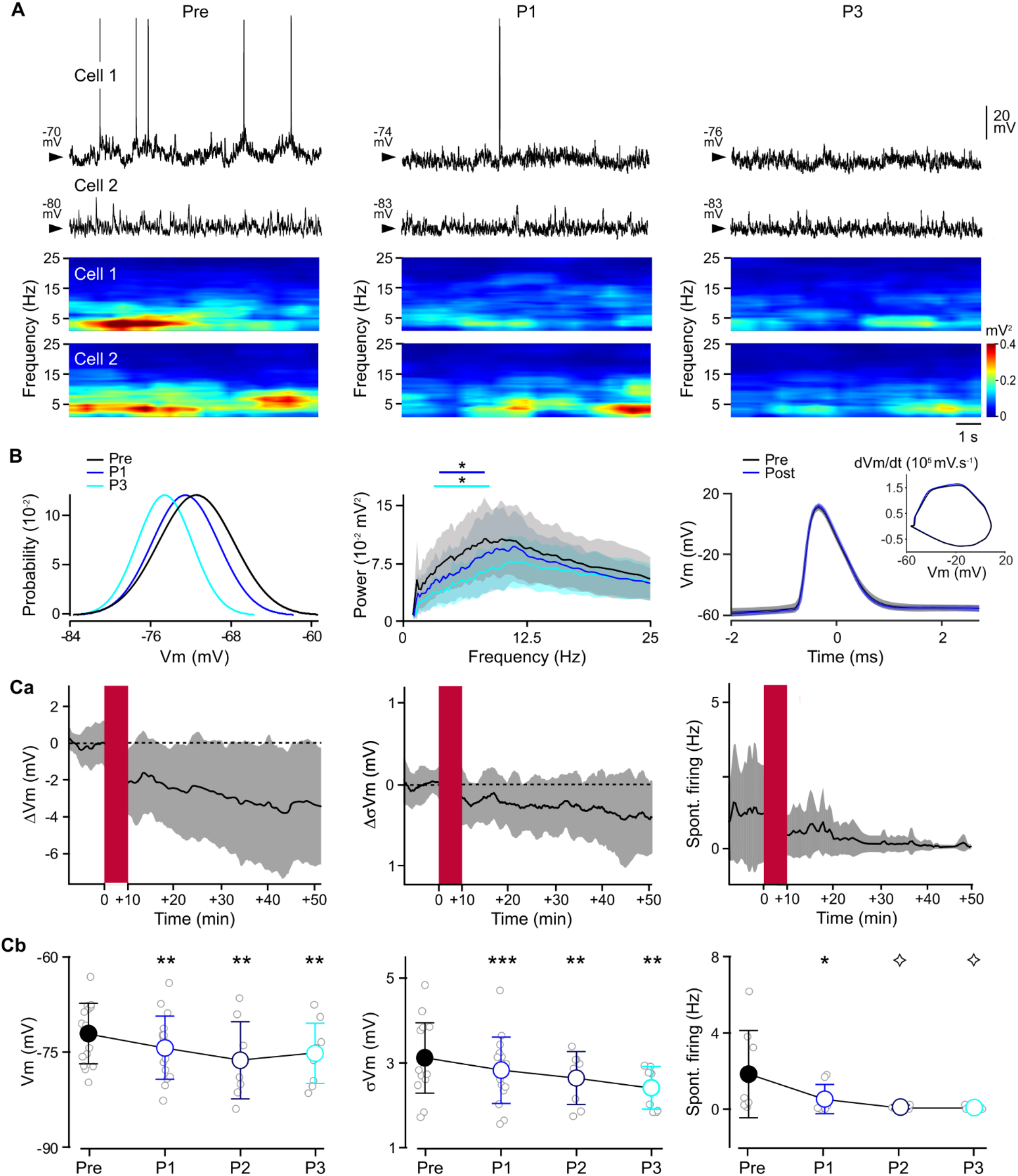
LI-rTMS reduces cortical neuron spontaneous activity. (**A**) Spontaneous intracellular activity recorded from active (cell 1) and silent (cell 2) S1 neurons before LI-rTMS (Pre) and during the first (P1) and third (P3) post-stimulation periods. The associated time-frequency maps, after removal and interpolation of APs, are temporally-aligned below the records. (**B**) Left: probability densities of Vm values computed from cell 2 during the different periods (60 sec of recording, bin size 1 mV). Middle: mean (solid lines) ± SD (shaded areas) frequency power spectra computed from intracellular subthreshold signals in the pre- and post-stimulation periods (*n* = 16 neurons) showing a progressive decrease in the power of the 5-10 Hz frequency band after LI-rTMS (Wilcoxon signed rank test with Benjamini-Hochberg’s post hoc correction, comparing P1 and P3 with Pre). Right: superimposed examples of average ± SD (shaded areas) of spontaneous APs and corresponding phase-plots (inset) during Pre (black trace, *n* = 138) and Post (blue trace, *n* = 28) periods. **(C)** Time-dependent changes in spontaneous activity after 10 minutes LI-rTMS. **(Ca)** Time course of the mean change (black line) ± SD (gray shaded area) in Vm (after subtraction of the mean Pre value), in the amplitude of Vm fluctuation (σVm; after subtraction of the mean Pre value), and in the spontaneous firing frequency of active neurons (Spont. Firing), calculated over successive 30-second periods before and after LI-rTMS application (red rectangle). **(Cb)** Comparison of Vm, σVm and spontaneous firing rate values (mean ± SD) during baseline (Pre) and at the three post-LI-rTMS times (P1, P2, P3) [RMANOVA and Bonferroni post-hoc comparisons; Vm: *n* = 16, 16, 8 and 9, F_0.39, 3.89_ = 12.35, *P* = 0.034; σVm: *n* = 16, 16, 8 and 9, F_1.36, 13.55_ = 11.69, *P* = 0.0025; spontaneous firing: *n* = 8, 8, 3 and 3, F_1.64, 6.03_ = 9.74, *P* = 0.015]. Here and in the following figures, the gray circles in the pooled-data graphs indicate values for individual neurons. In (Ca) and (Cb), analysis of ongoing firing was restricted to neurons which were spontaneously active during baseline. \**P* < 0.05, ** *P* < 0.01, *** *P* < 0.001, ns: non-significant; ◊ = too few data for post-hoc comparison.

In all tested neurons, 10 minutes continuous 10 Hz LI-rTMS induced membrane hyperpolarization compared to Pre (Fig. 2A-C), which was present immediately after LI-rTMS (P1: -2.3 ± 2.0 mV, *n*=16 neurons; *P*<0.01) and persisted until the end of recording (P3: -2.3 ± 1.9 mV, *n*=9 neurons; *P*<0.01) (Figs. 1Cb, 2A-C), resulting in a negative shift of the Vm distribution during the post-stimulation periods (Fig. 2B left). Background synaptic activity amplitude (Vm standard deviation) was also reduced after LI-rTMS in both active and silent neurons (Fig. 2A), reaching a 10.8 ± 12.2 % decrease at P3 as compared to the pre-stimulation period (*P*<0.01, *n*=9 neurons) (Fig. 2A-C). This was associated with reduced power of synaptic activity for frequencies below 10 Hz (Fig. 2A, B). This dampening of synaptic noise amplitude was unexpected because sustained Vm hyperpolarization should increase synaptic current driving force and therefore synaptic potentials, particularly given the lack of inward membrane rectification in the voltage-current (*V-I*) relationships in our recorded cells (fig. S1Bb).

Consistent with attenuated intracellular synaptic events and cell hyperpolarization, the spontaneous firing of these cortical neurons was markedly reduced after LI-rTMS. Five of the 8 neurons active prior to LI-rTMS became completely silent by P2, while firing of the 3 remaining cells decreased to 0.07 ± 0.13 Hz at P3 (Fig. 2A-C). This failure of cell firing was not due to altered ionic AP mechanisms, as their amplitude, half-width duration and threshold voltage did not change (Fig. 2B right, table S1); it more likely reflected the decreased background synaptic drive.

### LI-rTMS alters neuronal integrative properties and input-output relationship

We next tested if LI-rTMS altered the excitability of S1 pyramidal neurons by changing their integrative properties, *i*.*e*. their ability to process subthreshold membrane currents, and their input-output relationship before and after stimulation.

We first examined changes in membrane input resistance (Rm) and membrane time constant (τm), two parameters that govern, respectively, the ability of synaptic currents to change Vm and the efficacy with which a neuron summates synaptic potentials over time. Rm values, extrapolated from the slope of the *V-I* curves (fig. S1B), were already significantly reduced at P1 (compared to Pre: -14.1 ± 6.6 %, *n* =11 neurons; *P*<0.001), a decrease that persisted for up to 40 min after the end of stimulation (*n*=9 neurons; *P*<0.01) (Fig. 3A, C). The decrease in Rm was not caused by LI-rTMS-mediated changes in the morphology of recorded neurons, whose anatomical properties were comparable to those classically described for S1 pyramidal cells (Brecht et al., 2003; Mahon and Charpier, 2012; Manns et al., 2004) (Fig. 1B). τm values also showed a persistent decrease that closely matched the reduction in Rm (Fig. 3C). The attenuation of Rm and τm by LI-rTMS may contribute to the smaller amplitude of spontaneous synaptic potentials and their ineffective summation to elicit APs.

**Fig. 3.**
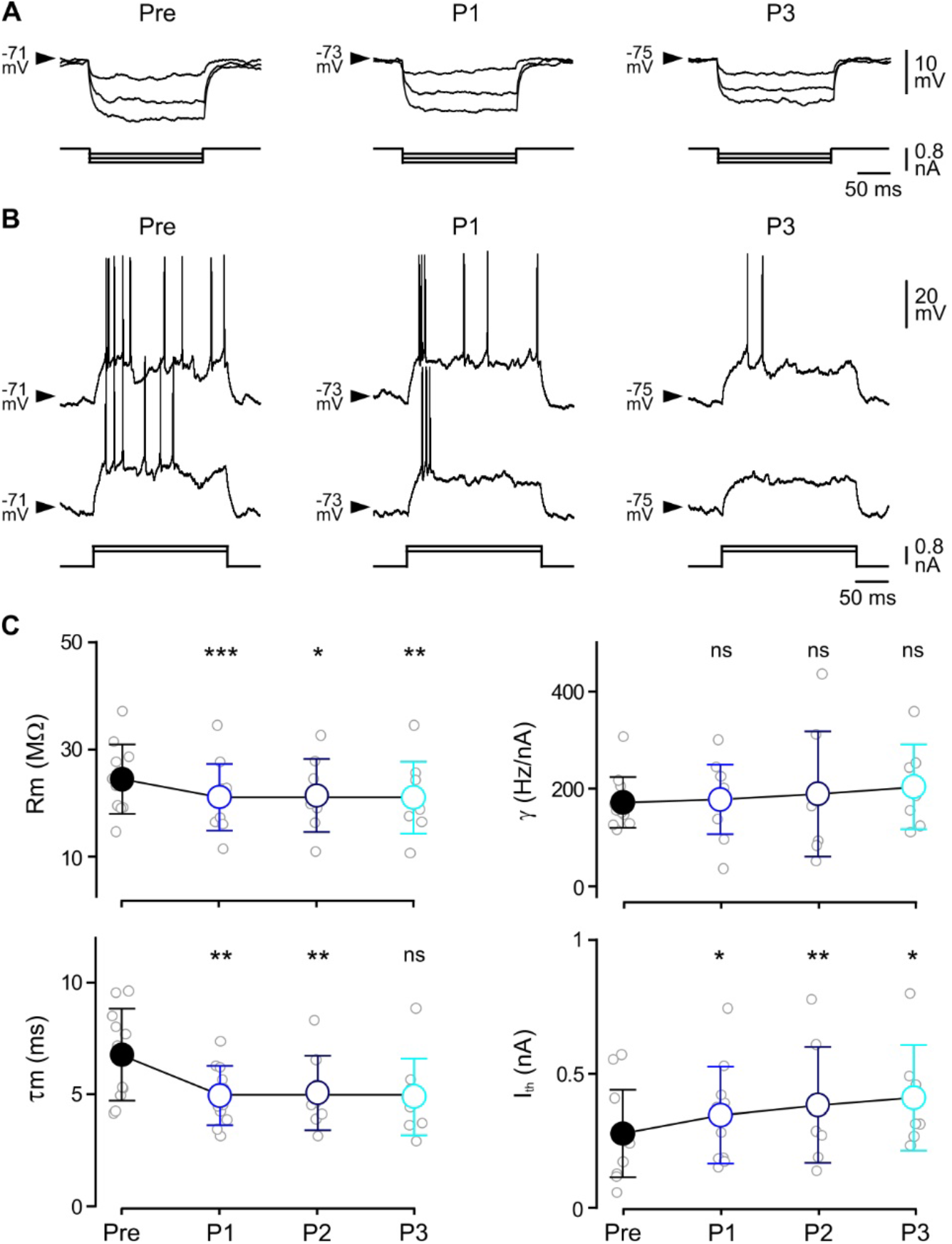
LI-rTMS negatively alters resting integrative properties and intrinsic excitability. (**A**) Average (*n* = 6–30 trials) voltage responses (upper traces) of an S1 pyramidal neuron to hyperpolarizing current steps of increasing intensity (−0.2 to -0.6 nA, lower traces) before (Pre) and after LI-rTMS (P1, P3). (**B**) Current-evoked firing responses of an S1 neuron to depolarizing current pulses of increasing intensity during Pre, P1 and P3. (**C**) Population data showing the temporal evolution of mean (± SD) values of Rm, τm, current threshold (I_th_) and neuronal gain (γ) values. [RMANOVA and Bonferroni post-hoc comparisons; Rm: *n* = 11, 11, 8 and 9, F_0.14, 1.19_ = 10.54, *P* = 0.08; τm: *n* = 11, 11, 8 and 9, F_1.61, 13.44_ = 8.24, *P* = 0.006; I_th_: *n* = 12, 11, 8 and 7, F_0.72, 5.55_ = 16.91, *P* = 0.01; γ: *n* = 12, 11, 8 and 7, F_0.14, 1.11_ = 0.66, ns]. \**P* < 0.05, ** *P* < 0.01, *** *P* < 0.001, ns: non-significant.

As changes in Vm and Rm can affect the reactivity of a neuron to its inputs (Altwegg-Boussac et al., 2014; Silver, 2010), we looked for possible LI-rTMS-induced modifications in the input-output relationship. We injected positive current pulses of increasing intensity during the different recording periods and counted the number of evoked APs, generating frequency-current (*F-I*) curves (fig S1Cb, Fig. 3B). After LI-rTMS, *F-I* curves progressively shifted to the right, leading to a ∼50% increase of the threshold current values by P3 (*n*=7 neurons; compared to Pre, *P*<0.05) (Fig. 3B, C). This reduced neuronal responsiveness to small excitatory inputs was accompanied by relatively stable neuronal gain, defined as the slope of the linear portion of *F-I* curves, throughout the post-stimulation period (*P* = 0.23) (Fig. 3C).

### Low-intensity magnetic pulses trigger APs

To identify the mechanisms initiating the long-lasting inhibitory effects of LI-rTMS on cortical activity and integrative properties, we investigated the immediate effects of LI-rTMS during the stimulation period. A practical difficulty arises from the brief (∼1.6 ms) large-amplitude artifacts generated by each LI-rTMS pulse (Figs. 1, 4, fig. S2), which obscures the early part of the neuronal response. However, we noticed in all recorded neurons (*n*=16) that the terminal part of many artifacts was rapidly followed by a biphasic voltage waveform resembling an intracellular AP (Fig. 4A, fig. S2A). We thus removed the stimulation artifacts off-line from traces displaying an apparent AP waveform (see Methods, and Figs. 4, 5 and fig. S2 for details). The artifact-free recordings revealed intracellular waveforms whose shape, peak amplitude and kinetics matched those of spontaneous APs recorded from the same neurons (Fig. 4Ab). The mean latency of AP-like events, calculated from the end of the stimulus to the peak of the waveform, was 0.6 ± 0.1 ms (*n*=16 neurons) and the percentage of individual magnetic pulses immediately preceding these waveforms was 24 ± 31 % (*n*=16 neurons; Fig 4B, bottom left), with a high cell-to-cell variability (from 0.3 to 99.8 %). Given that magnetic intensity decreases by distance-squared, we tested whether this inter-neuronal variability depended on the depth of recorded cells; there was no correlation (*r*^*2*^ = 0.03). Notably, these AP-like events are not non-biological artifacts resulting from direct action of the magnetic field on the recording electrode, as they did not occur in control experiments in which a model cell was used to simulate the recording of a pyramidal neuron (fig. S2B).

**Fig. 4.**
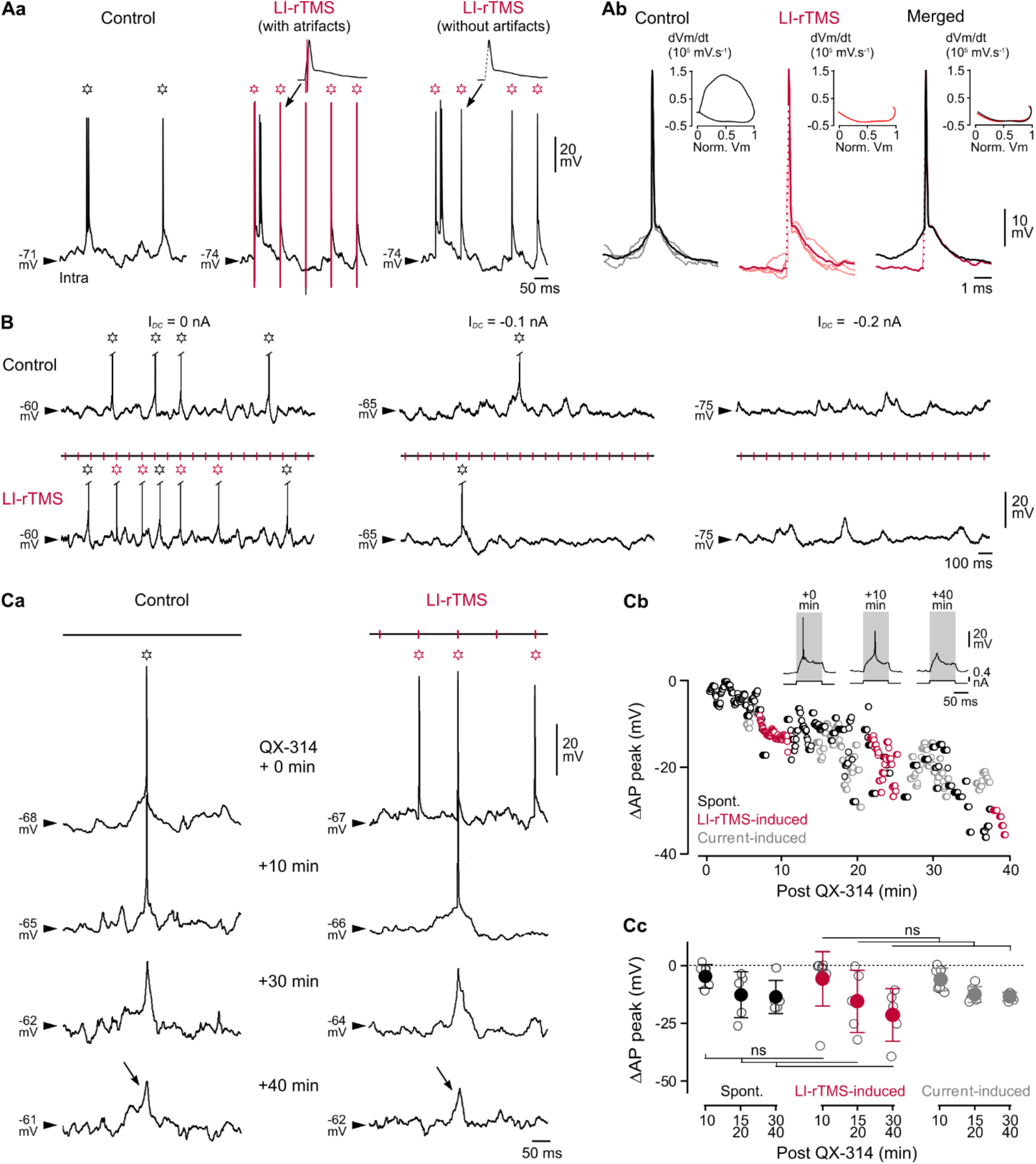
Evidence for LI-rTMS-induced APs. **(A)** Comparison of spontaneous and LI-rTMS-induced APs. **(Aa)** Intracellular activity of an S1 pyramidal neuron in control and during LI-rTMS, before (with artifacts, vertical red lines) and after (without artifacts) removal of stimulation artifacts. Here and in the following panels, open stars mark the occurrence of spontaneous (black) and putative LI-rTMS-evoked (red) APs. A typical example of a putative LI-rTMS-evoked AP, with and without artifact, is enlarged above the records. Note that the depolarizing phase of the LI-rTMS-evoked AP is hidden within the artifact. **(Ab)** *S*uperimposition of 3 spontaneous (black) or LI-rTMS-induced (red) APs (light-colored traces) with their corresponding average waveform (deeper-colored traces). The AP amplitude is normalized and LI-rTMS artifacts are indicated by dotted red lines. Note the similarity in the repolarization phase of spontaneous and LI-rTMS-induced APs when average waveforms are merged (right). The insets show the corresponding AP phase-plots (with only the repolarization phase illustrated for LI-rTMS induced APs). **(B)** Spontaneous and LI-rTMS-induced APs are blocked by membrane hyperpolarization. Cortical intracellular activity in absence of (Control, top traces) and during LI-rTMS (bottom traces) at rest (left panel) and during membrane hyperpolarization induced by intracellular injection of negative *DC*-current (middle and right panels). Red vertical markers above the traces indicate the position of the removed artifacts. **(C)** AP blockade by QX-314 in control and during LI-rTMS. **(Ca)** Examples of intracellular records obtained in absence of (left) and during (right) LI-rTMS at the indicated times after impalement with a microelectrode containing 100 mM QX-314. Times after impalement are indicated between each set of traces. Note the progressive decrease in amplitude of both spontaneous and LI-rTMS-triggered APs, which start to resemble sharp synaptic depolarizations at 30–40 min after onset of QX-314 diffusion (arrows). **(Cb)** Changes in peak potential value of individual spontaneous (black), LI-rTMS-induced (red) and current-induced (gray) APs in an S1 neuron, as a function of time after onset of QX-314 diffusion. The peak value of the first AP in each group, measured just after cell impalement, was taken as reference and artificially set to zero mV. Insets show representative current-evoked responses at the indicated times after cell impalement. **(Cc)** Pooled data comparing the peak potential decrease of spontaneous (black), LI-rTMS-induced (red) and current-induced (gray) APs at different times after QX-314 diffusion onset. [Mixed Effects ANOVA; Spont: *n* = 7, 5, 5 neurons; LI-rTMS-induced: *n* = 8, 5, 5 neurons; Current-induced: *n* = 8, 6, 5. Effect of time, F_1.21, 9,72_ = 16,64, *P* = 0.0017; Effect of stimulation, F_1.33, 10,64_ = 20.13 *P* = 0.0006). Between groups ANOVA: t0, F_2, 24_ = 2.34; t10, F_2, 20_ = 1.56; t15-20, F_2, 13_ = 0.68; t30-40, F_2, 12_ = 1.81. At each time-point, all comparisons, ns]. ns: non-significant.

**Fig. 5.**
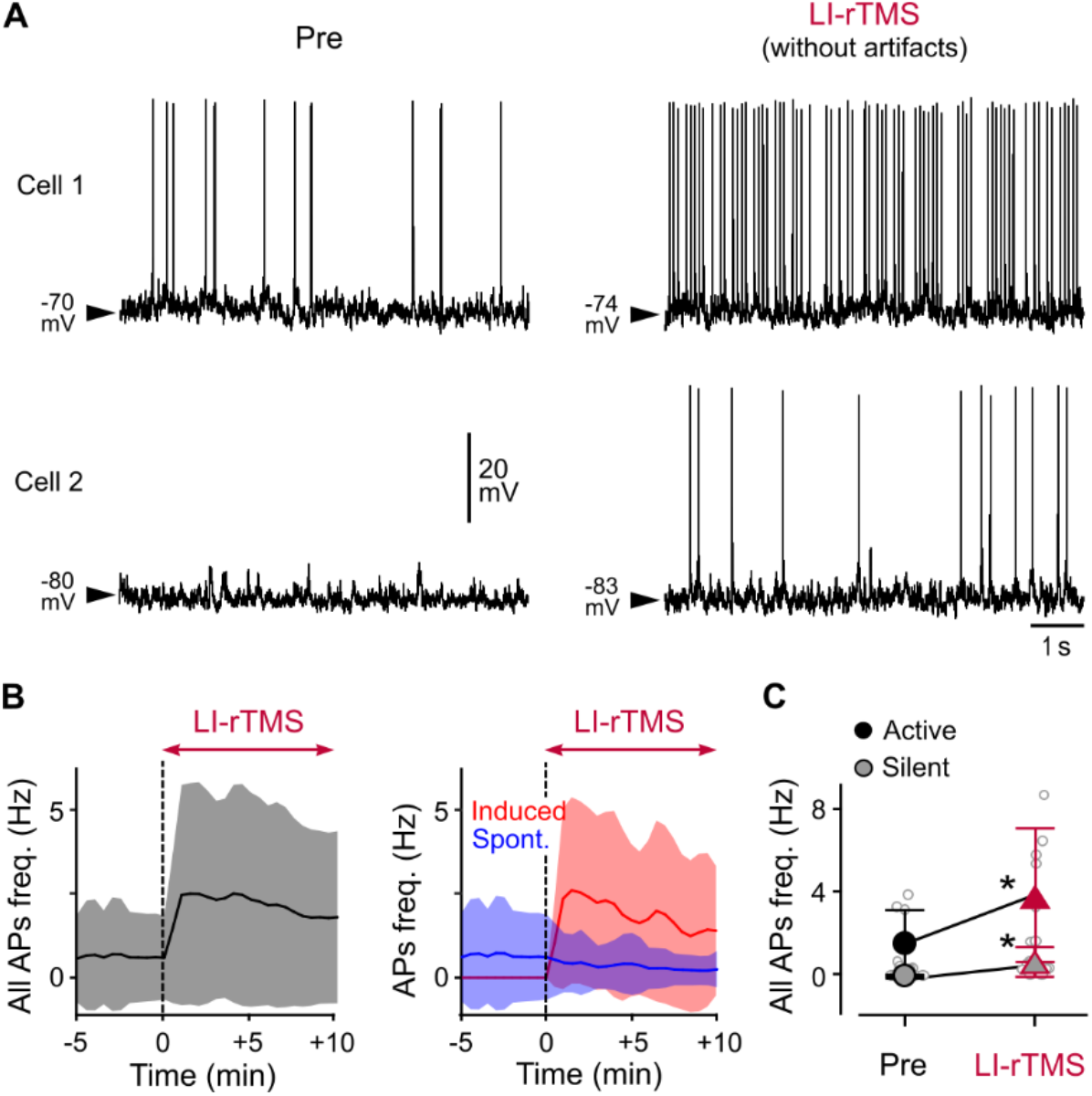
Changes in firing rate during LI-rTMS. **(A)** Intracellular activity of representative active (Cell 1, located in layer 2/3) and silent (Cell 2, located in layer 5) cortical neurons recorded before (Pre) and during LI-rTMS after artifact removal. **(B)** Time course of the mean (± SD, *n* = 16 neurons) changes in global firing frequency (including both spontaneous and LI-rTMS-induced APs; left) and of the specific rate of spontaneous (blue line) and TMS-mediated (red line) APs (right), calculated over 30-second periods before and during LI-rTMS (red arrow). **(C)** Summary plot comparing the mean (± SD) total AP frequency in Pre and LI-rTMS periods, separating cells that were active (black circles) and silent (gray circles) during baseline. [*n* = 8 active and 7 silent neurons, Wilcoxon Signed Rank test, \**P* < 0.05].

To confirm that the high-voltage waveforms were indeed APs, we tested their sensitivity to sustained Vm hyperpolarization - which prevents firing by moving the neuron’s Vm further from the AP threshold - or to the blockade of voltage-gated sodium channels by the lidocaine analog QX-314. In the absence of LI-rTMS, spontaneous firing frequency which was 1.8 ± 1.6 Hz at rest (Vm = -71.9 ± 7.0 mV, *n*=6 tested neurons), declined with imposed *DC* hyperpolarization (from -0.1 to -2 nA), and fell to zero when Vm reached -81.2 ± 8.1 mV (*n*=6 neurons) (Fig. 4B, top). Vm hyperpolarization was associated with increased amplitude of synaptic fluctuations, as expected from a greater driving force of excitatory synaptic currents (Fig. 4B, top right). Hyperpolarizing these same cortical neurons during short periods of LI-rTMS similarly decreased the rate of the AP-like events, from 4.0 ± 2.7 Hz at rest (*n*=6 neurons) to a residual rate of 0.04 ± 0.05 Hz at the most negative membrane polarization tested (Vm = -81.2 ± 8.1 mV, *n*=6 neurons) (Fig. 4B, bottom).

Second, to test whether the putative LI-rTMS-induced APs had the same ionic mechanisms as spontaneous APs, we perfused the intracellularly recorded neurons with QX-314 (*n*=9) and compared the drug-related changes in amplitude of both types of AP. In each neuron, measurements were made during alternating control and stimulation periods of 5–160 seconds. The peak potential of spontaneous APs was progressively reduced (*P*<0.05) by the QX314 (Fig. 4C) and this time-dependent reduction was accompanied by a mean Vm depolarization of 5–10 mV, as classically observed in QX-314-filled cortical neurons (Benardo et al., 1982; Connors and Prince, 1982). The decrease in peak values of LI-rTMS-induced waveforms and spontaneous APs was almost identical for any given time after the onset of QX-314 diffusion (*P*>0.2 for each recording time), reaching ∼ -14–20 mV in the neurons that could be recorded until 30–40 min after cell impalement (*n*=5) (Fig. 4C). Because the QX314-reduced APs were occasionally difficult to distinguish from spontaneous depolarized synaptic potentials in recordings of long duration (Fig. 4Ca, arrows), we also examined the effects of QX-314 on APs evoked by depolarizing current steps (0.2–0.4 nA, *n*=9 neurons) (Fig. 4Cb, Cc). Again, the time-course of the reduction in current-induced AP peak potentials was similar to that measured for LI-rTMS-induced APs (*P*>0.1 for each recording time) (Fig. 4Cb, Cc).

The LI-rTMS-induced APs occurred in all recorded cortical neurons, regardless of their firing frequency prior to stimulation (Fig. 5A-C). This led to a significant increase in the overall firing rate (including both spontaneous and LI-rTMS-evoked APs) throughout the LI-rTMS period (*n*=16 neurons; *P*<0.05) (Fig. 5B). However, the rate of spontaneous APs, when measured separately, was decreased by almost 45% during LI-rTMS (Pre: 1.8 ± 2.3 Hz, *n*=8 active neurons, LI-rTMS: 0.6 ± 0.8 Hz, *n*=8 neurons; *P* < 0.01) (Fig. 5B). The induced APs did not result from a LI-rTMS-mediated increase in synaptic activity or sustained membrane depolarization since artifact-free records did not show significant change in the amplitude of Vm fluctuations in association with the firing induced by magnetic pulses (Figs. 4, 5, and fig. S2A). However, their short latency and ability to be evoked from relatively hyperpolarized membrane potentials indicate that LI-rTMS-evoked APs probably resulted from activation of an axonal site located close to the soma.

These results demonstrate for the first time that 10 Hz LI-rTMS can elicit AP firing in pyramidal neurons, with kinetic and ionic mechanisms similar to spontaneous APs, despite inducing an electric field estimated to be at least 100 times lower than that used in high-intensity rTMS. They also demonstrate that 10 minutes continuous 10 Hz LI-rTMS produces robust and long-lasting inhibitory effects on the activity and integrative properties of individual pyramidal neurons.

### Action potentials evoked by intracellular current injection do not reproduce LI-rTMS effects

In order to determine whether the LI-rTMS-induced firing was responsible for the changes in cortical neuron activity and responsiveness, we examined, in another group of S1 neurons, the impact of an AP discharge evoked by intracellular injection of current steps. Single cortical neurons (*n*=7) were challenged with repetitive depolarizing current pulse stimulation (rCPS) delivered intracellularly at10 Hz for 10 min (Fig. 6A). Intensity of current steps was adjusted in each neuron to obtain a proportion of suprathreshold stimulations (∼30%) and an overall firing frequency comparable to those produced by LI-rTMS (5.9 ± 4.6 Hz, *n*=7 neurons, including both spontaneous and evoked APs; *P*>0.4) (Fig. 6B). The effects of rCPS were compared to a subgroup of magnetically stimulated neurons (*n*=9) with similar baseline membrane electrical properties, including mean Vm, spontaneous firing frequency and Rm (*n*=9; *P*>0.1 for each parameter).

**Fig. 6.**
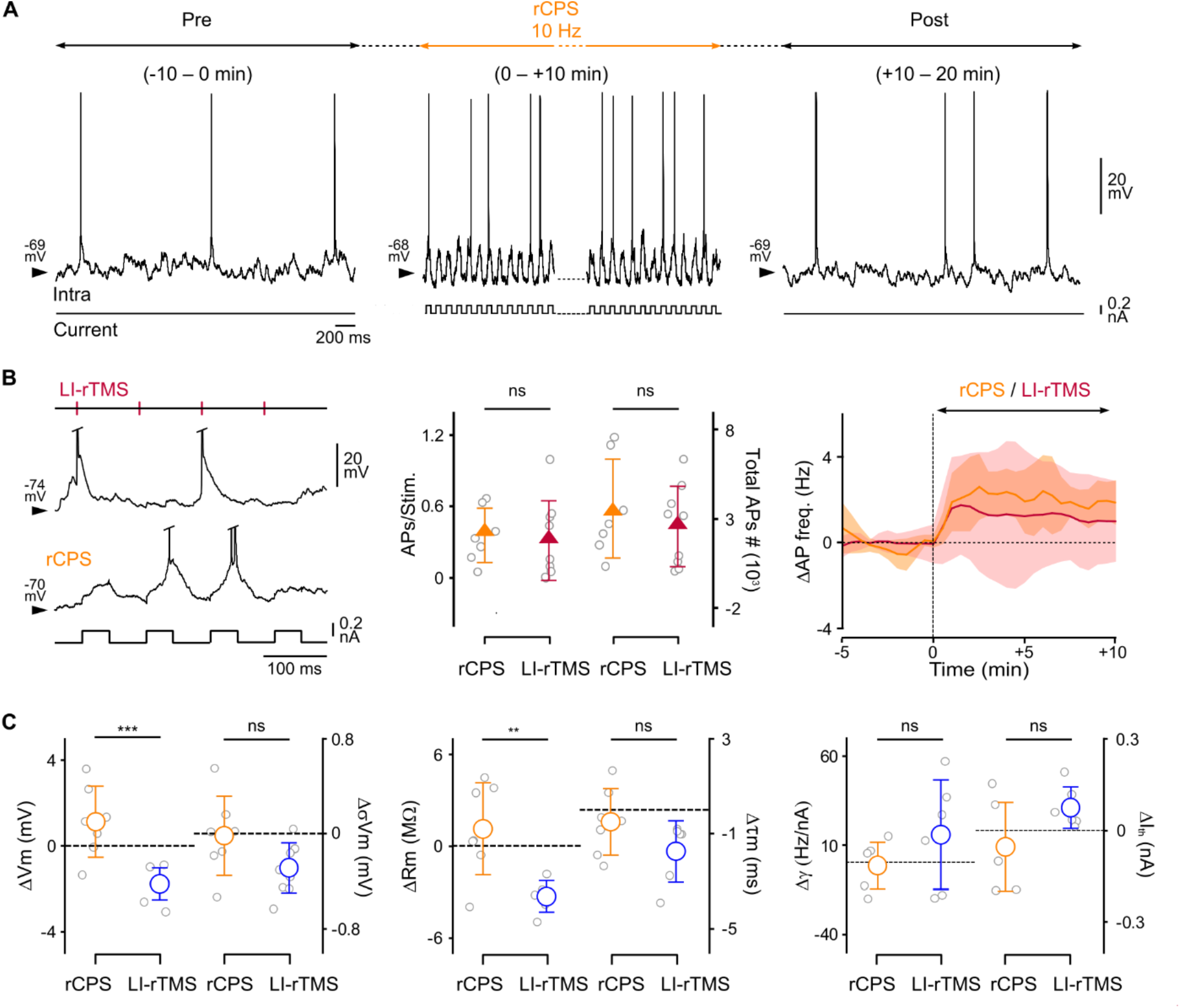
Repetitive current-evoked firing in a single cortical neuron does not reproduce LI-rTMS effects. **(A)** Cortical intracellular activity before, during and after application of repetitive current pulse stimulation (rCPS). The rCPS protocol consisted of injecting a series of depolarizing current pulses (0.2 nA, 40 ms duration) at 10 Hz for 10 min (lower trace). **(B)** Comparison of neuronal firing during rCPS and LI-rTMS protocols. Left: Short periods of intracellular activity during LI-rTMS (top trace) and rCPS (bottom trace) paradigms. Onset times of removed LI-rTMS artifacts are indicated by red vertical markers and the timing of depolarizing current pulses is shown below the bottom trace. Middle: Pooled data comparing the mean (± SD) number of APs/stim. evoked by LI-rTMS or current pulses, and the total number of APs (evoked and spontaneous) measured over the whole stimulation period in response to rCPS (*n* = 7 neurons, orange triangles) or LI-rTMS (*n* = 9 neurons, red triangles). Right: Time course of the mean change (solid lines, *n* = 7 neurons) ± SD (shaded areas) in AP frequency (after subtraction of the mean Pre value) calculated over successive 30-second periods, before and during rCPS (orange) or LI-rTMS (red) protocols. Dotted line indicates the mean Pre value, artificially set to zero. **(C)** Summary data comparing the changes in Vm, σVm, Rm, τm, γ and Ith values between pre- and post-stimulation periods after application of rCPS (orange circles) or LI-rTMS (blue circles) protocols. Dashed lines indicate the mean pre-stimulation value (Pre), artificially set to zero [Vm: *n* = 7, 8; σVm: *n* = 7, 9, Rm: *n* = 7, 6; γ: *n* = 7, 6; Ith: *n* = 7, 6; unpaired Student t-test; and τm: *n* = 7, 6; Mann Whitney Rank Sum test; ** *P* < 0.01, *** *P* < 0.001, ns: non-significant].

The first obvious difference between effects of current-evoked APs (rCPS) and LI-rTMS was that rCPS did not produce neuronal hyperpolarization during the post-stimulation period (Fig. 6C, left and fig. S3D). This was accompanied by stable Vm fluctuations and spontaneous firing frequency (Pre *versus* post-rCPS: *P*>0.2 for each parameter) (fig. S3A, D). Furthermore, comparison of *V-I* curves before and after rCPS showed no significant modification of Rm (*n*=6 neurons; Pre, 17.2 ± 6.9 MΩ *vs*. Post, 17.9 ± 7.5 MΩ; *P* = 0.14) or τm (*n*=6 neurons; Pre, 6.5 ± 1.8 ms, vs. Post, 6.3 ± 1.6 ms; *P* = 0.28) (Fig. 6C, middle and fig. S3B, D). Finally, analysis of *F-I* curves revealed heterogeneous modification in the sensitivity of cortical neurons to weak depolarizing inputs. Although average current threshold values did not significantly change (*P*=0.4), most neurons (*n*=4 out of 6 tested cells) had a significantly decreased firing threshold (ΔI_th_ = -0.14 ± 0.07 nA, *n*=4; *P*<0.05) following rCPS. This modulation of cortical responsiveness was not accompanied by modification in the neuronal gain (*P*>0.4) (Fig. 6C right and fig. S3C, D). Thus, the firing of single cortical neurons at a similar frequency to that induced by magnetic stimulation (∼ 5 Hz) is not sufficient to reproduce, on its own, the LI-rTMS-related changes in neuronal excitability.

## Discussion

To our knowledge, this study is the first to investigate the immediate (online) and after- (offline) effects of low-intensity rTMS on the electrical properties of single neurons *in vivo*. Because rTMS-induced cortical responses depend on the ongoing brain state (Gersner et al., 2011; Silvanto and Pascual-Leone, 2008), we used experimental conditions which retain stable background cortical activity, similar to the waking state, during the long intracellular recordings (Altwegg-Boussac et al., 2014; Bruno and Sakmann, 2006; Constantinople and Bruno, 2011). Our results indicate that 10 minutes continuous 10 Hz LI-rTMS alters the passive membrane properties of S1 pyramidal neurons, reduces their intrinsic excitability and attenuates their spontaneous firing for at least 40 min. Moreover, low-intensity magnetic pulses triggered APs in all recorded neurons. These findings deepen our knowledge on the cellular mechanisms underlying LI-rTMS-induced plasticity and may guide the development of new stimulation patterns for clinical treatment.

### LI-rTMS induces action potentials in stimulated neurons

A major and unexpected finding of our study was that low intensity (10 mT) magnetic pulses elicited APs in all recorded neurons (25% of pulses on average). This challenges the common assumption that LI-rTMS only acts through subthreshold mechanisms because it induces electrical fields weaker than those required for neuronal activation (Chan and Nicholson, 1986; Grehl et al., 2016; Radman et al., 2009). Our result is however coherent with weak electric fields inducing short-latency spiking in retinal ganglion cells *in vitro* (Bonmassar et al., 2012) and medium-intensity (60 mT) magnetic pulses transiently activating sodium currents to facilitate AP firing in S1 pyramidal neurons (Banerjee et al., 2017) in slices. The percentage of magnetic pulses initiating APs varied between cells, consistent with variable pyramidal neuron sensitivity to magnetic and electric fields, depending on their location, morphology and intrinsic excitability (Radman et al., 2009; Rotem and Moses, 2008) as well as fluctuations in their spontaneous firing (Pasley et al., 2009).

The mechanisms underlying magnetic pulse-induced neuronal activation remain unclear. In high-intensity TMS, trans-synaptic or direct neuronal depolarization are thought to initiate APs in the axon (Müller-Dahlhaus and Vlachos, 2013; Pashut et al., 2014; and references therein). Here, LI-rTMS-induced APs had short latency, were not associated with increased subthreshold synaptic events, and could be triggered from a hyperpolarized Vm, suggesting direct activation of the axon initial segment, whose high concentration of voltage-gated sodium channels confers a lower AP threshold (Eccles, 2013; Kole et al., 2008).

### Continuous 10 Hz LI-rTMS durably decreases cortical excitability and activity

The main after-effects of LI-rTMS included smaller amplitude background synaptic noise, substantial membrane hyperpolarization and marked reduction or cessation of spontaneous firing. These changes in spontaneous activity were accompanied by attenuated neuronal integrative properties (correlated decrease in Rm and τm) and reduced intrinsic excitability (higher minimal current firing threshold).

The likely primary origin of these effects, whether altered synaptic events or intrinsic cellular properties, can be deduced from our data. Long-term depression of synaptic inputs onto the recorded neurons, which could explain the observed reduction in synaptic activity, would decrease global membrane conductance (due to smaller excitatory synaptic conductance) and thus increase Rm and τm values (Destexhe et al., 2003) – the opposite of our findings. Rather, we suggest that LI-rTMS-mediated inhibition of neuronal activity originates from alteration to cortical neuron integrative properties. Indeed, the initial reduction in Rm and τm, which attenuates neuronal responsiveness to their synaptic inputs, would explain both the reduced amplitude of Vm fluctuations and membrane hyperpolarization. Given the inverse correlation linking current firing threshold to Vm in cortical neurons (Altwegg-Boussac et al., 2014; Carton-Leclercq et al., 2021), the membrane hyperpolarization would in turn explain the rightward shift of input-output relations and the reduced intrinsic excitability. Finally, all these cellular modifications will combine to reduce the efficiency with which cortical neurons process synaptic currents and generate APs and, thus, explain the progressive decrease of spontaneous firing. This reduced firing in individual neurons will contribute to global reduction of synaptic activity flow within the stimulated cortical network, which will further reduce synaptic and firing activity. Furthermore, increased activity of neocortical GABAergic interneurons could also contribute to reducing pyramidal cell activity, by producing a robust shunting effect on pyramidal neuronal membrane (Tremblay et al., 2016) and lowering Rm (due to larger inhibitory synaptic conductance).

The precise subcellular mechanisms through which LI-rTMS exerts its effects warrant further investigation. We hypothesize that increased neuronal firing during stimulation opens voltage-gated calcium channels, increasing intracellular calcium and activating calcium-dependent enzymatic cascades that regulate ionic conductances (Delord et al., 2007; Xu and Kang, 2005). In line with this hypothesis, pyramidal neuron firing induced by medium-intensity (60 mT) rMS *in vitro* is associated with increased cytosolic calcium (Banerjee et al., 2017). Moreover, our data showing changes in Rm and current firing threshold together with stable neuronal gain after LI-rTMS suggest that the long-lasting decrease in neuronal excitability is due to modulation of potassium currents activated around resting Vm (Naudé et al., 2012). This suggestion is supported by previous reports, in which similar depression of intrinsic cell excitability was attributed to calcium-dependent upregulation of slowly-inactivating potassium currents (Li et al., 2004; Lombardo et al., 2018).

Our results also indicate that induced APs, and their associated changes in calcium concentration, may not be sufficient to trigger the time-dependent post-LI-rTMS effects. Simply replicating the level of LI-rTMS-induced AP firing by repeated intracellular injection of current pulses in single neurons did not significantly modify resting and active membrane properties. This suggests that electric field-induction is only one of several mechanisms activated by electromagnetic pulses. We have previously shown that cryptochrome magnetoreceptors are required for neuroplasticity effects of LI-rTMS (Dufor et al., 2019) and modulate reactive oxygen species (ROS) (Sherrard et al., 2018). In cell redox physiology, ROS activate ryanodine receptors to release calcium from intracellular stores (Görlach et al., 2015), a phenomenon we have demonstrated in cortical neurons in response to LI-rTMS (Grehl et al., 2015). It is therefore likely that both voltage-gated calcium channel activation and calcium release from intracellular stores are required for the long-term modulation of pyramidal neuron activity.

### Relevance of LI-rTMS to human rTMS treatment

The LI-rTMS-induced inhibitory effects on neuronal excitability and firing is surprising in light of former studies using high-intensity rTMS. According to these reports, high-frequency magnetic stimulation (≥ 5 Hz) should increase cortical excitability (Bhattacharya et al., 2021; Cirillo et al., 2017; Huang et al., 2005), potentiate excitatory synaptic transmission and/or upregulate the expression of neurochemical markers of synaptic plasticity (Cirillo et al., 2017; Gersner et al., 2011; Lenz et al., 2015; Vlachos et al., 2012).

However, there are important differences between those reports and our study, which can explain the different outcomes. First, in some of these studies, rTMS effects on synaptic function were examined *ex vivo* after repeated daily sessions of high-frequency rMS (Cirillo et al., 2017; Gersner et al., 2011), which is not the case here. Second, in studies where acute effects of magnetic stimulation were measured, rMS was usually delivered directly to a brain slice or organotypic culture and not to an intact brain as in our *in vivo* experimental condition (Lenz et al., 2015; Tang et al., 2016; Vlachos et al., 2012). For example, in mouse S1 slices, 3 minutes of medium-intensity (85 mT) intermittent theta-burst magnetic stimulation reduced AP voltage threshold in pyramidal neurons and increased their current-evoked firing, without changing resting membrane properties (Tang et al., 2016). Discordance between results obtained *in vitro* and in living brains has already been reported: opposite changes in synaptic efficacy can be induced *in vivo* and *in vitro* at the same synapses by the same stimulation paradigm (Mahon et al., 2004). Thus differences between previous observations and our current *in vivo* findings reinforce that rTMS outcomes depend on ongoing activity in the targeted neural networks during the stimulation (Gersner et al., 2011; Silvanto and Pascual-Leone, 2008). Finally, most previous studies used magnetic fields at least ten times stronger than the one used in our study. High-intensity, high-frequency stimulation protocols require regular pauses to allow the coil to cool – resulting in short periods (a few seconds) of stimulation separated by pauses without stimulation (see Bhattacharya et al., 2021; Cirillo et al., 2017 for reviews), an intermittent pattern in contrast to our continuous 10 Hz. Theta-burst stimulation, for example, induces opposite effects if applied continuously versus intermittently (Huang et al., 2005). Thus, LI-rTMS effects on cell excitability are likely to depend on stimulation parameters, similar to the pattern-dependence of LI-rTMS effects on structural and biochemical plasticity (Dufor et al., 2019; Grehl et al., 2015).

Although the functional consequences of LI-rTMS-mediated depression of cortical excitability for S1 network function and behavior remain unknown, our LI-rTMS protocol could be useful in epileptic disorders, which are often associated with cortical hyperexcitability and paroxysmal network synchronization (Kimiskidis et al., 2014). An understanding of the immediate and lasting effects on neuronal activity, measured in real time, is an important advance for the development and evaluation of stimulation patterns in future clinical treatments.

In summary, our findings that even weak magnetic fields can activate neurons and modulate their excitability, change our understanding of low-intensity magnetic brain stimulation. We show that low-intensity approaches have some underlying processes in common with already established high-intensity clinical treatments, but without the constraints of special high-voltage equipment. This knowledge not only expands the range of stimulation protocols (pulse frequency and patterns), and therefore future potential clinical treatments, but also renders LI-rTMS treatments available beyond the hospital setting.

## Materials and Methods

### Animals, their preparation and surgery

All animal experiments received approval (APAFIS no. 18003-2019051019017280) from the *Charles Darwin* Animal Experimentation Ethics Committee (C2EA-05), in accordance with the guidelines of the European Union (directive 2010/603/EU). Experiments were conducted *in vivo* on adult Sprague Dawley rats (*n* = 31) of either sex (Charles River Laboratories, France). Under isoflurane (3.5%; Centravet, France) anesthesia, animals underwent tracheotomy for artificial ventilation (room air, 85 breaths.min^-1^, 2.6 ml.cycle^-1^) under neuromuscular blockade (gallamine triethiodide, 40 mg/2h i.m; Sigma-Aldrich, France). They were placed in a stereotaxic frame and small craniotomies were made above the left barrel field of S1 (1.2–1.3 mm posterior to bregma, 4.1–6.5 mm lateral to midline; Paxinos and Watson, 1986) to allow simultaneous recording of ECoG and intracellular activity of subjacent pyramidal neurons. After craniotomy, isoflurane was discontinued, but sedation and analgesia were maintained by repeated injection of sufentanil (3 µg/kg every 30 min, i.p.; Piramal) and local infiltration of lidocaine (2%, Centravet) in incision and pressure points. Sedation with sufentanil (Niemegeers et al., 1976) was chosen because it induces a globally stable background ECoG activity characterized by low amplitude desynchronized cortical waves resembling those encountered during the waking state (Altwegg-Boussac et al., 2017, 2014; Bruno and Sakmann, 2006; Constantinople and Bruno, 2011; Depaulis et al., 2016). It thus avoids anesthesia-related modulation of cortical responses to magnetic stimulation, as observed in both animal and human studies (Gersner et al., 2011; Silvanto and Pascual-Leone, 2008). Physiological parameters were continuously monitored during recordings (Altwegg-Boussac et al., 2017; Schramm et al., 2020). Each animal formed its own control, with LI-rTMS effects compared to baseline measurements obtained before the magnetic stimulation. After the recordings, animals were euthanized with euthasol (40%, i.p.; Centravet) for histological examination (see below).

### *In vivo* multi-scale electrophysiological recordings

Intracellular recordings were performed with glass micropipettes filled with 2-M potassium acetate (50-90 MΩ). Current-clamp recordings were amplified using an Axoclamp 900A amplifier (Molecular Devices, Union City, CA, USA) operating in bridge mode, filtered between *DC* and 30 kHz and digitized at 10 kHz (CED 1401plus, Spike2 software version 7.06; Cambridge Electronic Design). Intracellularly recorded neurons (*n* = 41), identified as pyramidal cells by their morphological and electrophysiological characteristics (Steriade, 2004), were located in layers 2/3 and 5 at depths ranging between 800–3300 µm below the cortical surface (Paxinos and Watson, 1986; Wilent and Contreras, 2004). In some experiments (*n* = 8), we added to the pipette solution 50–100 mM of QX-314 (Tocris Biosciences), a voltage-gated sodium channels blocker known to suppress APs in a time-dependent manner (Connors and Prince, 1982; Wilson and Kawaguchi, 1996).

Spontaneous ECoG activity was recorded with a low-impedance (∼60 kΩ) silver electrode placed on the dura above S1 and a reference electrode placed on a contralateral temporal muscle. Surface cortical signals were amplified using a differential *AC* amplifier (Model 1700; A-M Systems), filtered at 1 Hz– 1 kHz, and digitized at 3 kHz.

### Stimulation protocols

Continuous 10Hz LI-rTMS was applied for 10 minutes (6000 pulses) using a miniaturized round coil connected to an electromagnetic pulse generator (E-Lab5, Global Energy Medicine, Perth, Australia), which generated 300-μs monophasic pulses of 10mT intensity, as previously described (Makowiecki et al., 2014; Rodger et al., 2012). The coil was secured to a micromanipulator and placed above S1 (Fig. 1A). Brain-to-coil distance was minimized (∼2 mm) while assuring contact-free stimulation perpendicular to the cortical surface. We verified that LI-rTMS caused no electromagnetic interference in the recording leads and electrodes, with negative control experiments in which a model cell (CLAMP-1U Model Cell, Molecular Devices, Union City, USA) in ‘cell mode’ was attached to the headstage amplifier (fig. S2B).

Repetitive current pulse stimulation consisted of intracellular injections of depolarizing square current pulses (40 ms of duration) at 10 Hz for 10 min. The intensity of the current pulses (0.1–0.5 nA) was adjusted for each neuron to evoke an AP every 3–5 current pulses, mimicking the frequency of LI-rTMS-induced APs.

### Intracellular and ECoG signal analysis

Spontaneous activity, membrane excitability and integrative properties of cortical neurons were assessed before and after LI-rTMS. Average membrane potential (Vm) was calculated as the mean of the distribution of spontaneous subthreshold membrane potential values and the magnitude of Vm fluctuations was quantified as the standard deviation (σ) of the distribution (σVm). Time-frequency maps illustrating the spectral content of synaptic activity over time were constructed using a multitaper decomposition analysis with a sliding time window of 3 seconds (Oostenveld et al., 2011), after removing APs which were replaced by a piecewise cubic hermite polynomial interpolation. Power spectra were computed by averaging the time-frequency decomposition over a 10-minute period.

Mean spontaneous firing rate and Vm values were measured from recordings of at least 1 min. The regularity of neuronal firing was calculated by the CV2 method: here CV2 = (2|Δt_i+1_ - Δt_i_|)/(Δt_i+1_+ Δt_i_), and Δt_i_ = t_i_ – t_i-1_ is the inter-spike-interval (Altwegg-Boussac et al., 2014). AP properties were measured from spontaneously occurring APs. AP voltage threshold was defined as the Vm at which dVm.dt^-1^ first exceeds 10 mV.ms^-1^ (Mahon and Charpier, 2012). AP amplitude was calculated from threshold to peak potential and AP half-width was the duration of the waveform at half-amplitude.

Dynamic changes of Vm, σVm and spontaneous firing frequency were assessed over successive windows of 30 seconds. Recording periods during which cortical excitability was assessed (see below) were removed from this analysis and replaced by a linear interpolation.

Rm was calculated as the slope of the linear portion of voltage-current (*V-I*) plots generated following repeated injections of hyperpolarizing current of increasing intensity (200 ms at 2 Hz, 10–20 trials per intensity, -0.2 to -0.8 nA) or from the average voltage deflection induced by hyperpolarizing current pulses of -0.4 nA (200 ms, ≥ 10 trials). τm was derived from an exponential decay fit applied to the initial phase of the mean current-induced hyperpolarization. Neuronal transfer function (*F*–*I* relationship), which describes the cell output firing as a function of injected depolarizing current pulses of increasing intensity (200 ms at 2 Hz, 10–20 trials per intensity, 0.1–1.5 nA), was quantified before and after LI-rTMS. Linear regression of *F*–*I* curves yielded the threshold current for AP generation (I_th_; *x*-intercept) and neuronal gain (γ; slope) (Altwegg-Boussac et al., 2014; Mahon and Charpier, 2012).

### LI-rTMS artifact removal

To assess changes in cortical activity during the 10 minutes of LI-rTMS, we removed brief high-amplitude electrical artifacts induced by the magnetic pulses. We removed the recorded signal for 1.6 ms following each pulse onset and preserved continuity by linear interpolation (fig. S2), as described for human stimulation experiments (Amengual et al., 2017; Stengel et al., 2021). We validated this approach, by adding a new channel containing artificially generated stimulation events at 10 Hz for 10 minutes to control recordings. These false artifacts were then removed with the same procedure and the resulting electrophysiological signal compared to the original to ensure their similarity (Amengual et al., 2017).

### Neuronal labelling

Neurons were filled with neurobiotin (1.5% in the pipette solution; Vector Laboratories, Burlingame, USA) through repeated depolarizing current pulse injections over 10 minutes after the end of the post-stimulation period, as previously described (Mahon and Charpier, 2012). Rats were euthanized with euthasol and transcardially perfused with 0.3% glutaraldehyde-4% paraformaldehyde in PBS (0.1 M, pH 7.4; VWR Chemicals, France), and the neurobiotin-filled neurons revealed as previously described (Williams et al., 2016).

### Statistical analysis

Data were explored for normality, outliers and statistical test assumptions in Prism version 9.00 (GraphPad Inc., San Diego, USA). Group differences were assessed using paired two-tailed Student’s t-test, one-way Repeated Measures ANOVA, or non-parametric Wilcoxon signed rank test and the Mann-Whitney rank sum test. The Benjamini-Hochberg test was used to correct for multiple comparisons (Benjamini and Hochberg, 1995). Values are expressed as means ± SD and significance level was *P* < 0.05.

## Acknowledgments

The authors thank Sarah Lecas for histological processing and all the technical staff from the Histomics core facility of the Paris Brain Institute. Rats were housed in the PHENO-ICMice core facility, which is supported by 2 Investissements d’Avenir (ANR-10-IAIHU-06 and ANR-11-INBS-0011-NeurATRIS) and the Fondation pour la Recherche Médicale.

## Funding

Agence nationale de la recherche, ANR-19-CE37-0021

Agence nationale de la recherche, ANR-10-IAIHU-06

Agence nationale de la recherche, ANR-16-CE37-0021

Fondation Recherche Alzheimer, (MB scholarship)

## Author contributions

Conceptualization: SM, RS, SC, AL, AVC

Methodology: MB, SM, SC, RS, AL, AVC

Investigation: MB

Data analysis: MB, PB, CS, SM, RS

Supervision: SM, RS, SC, AL, AVC

Writing – original draft: SM, SC, RS, AL

Writing – review & editing: SM, SC, RS, AL, MB, PB, AVC, CS

## Data and materials availability

All data are available in the main text or the supplementary materials.

## Supplementary Materials

**Fig. S1.**
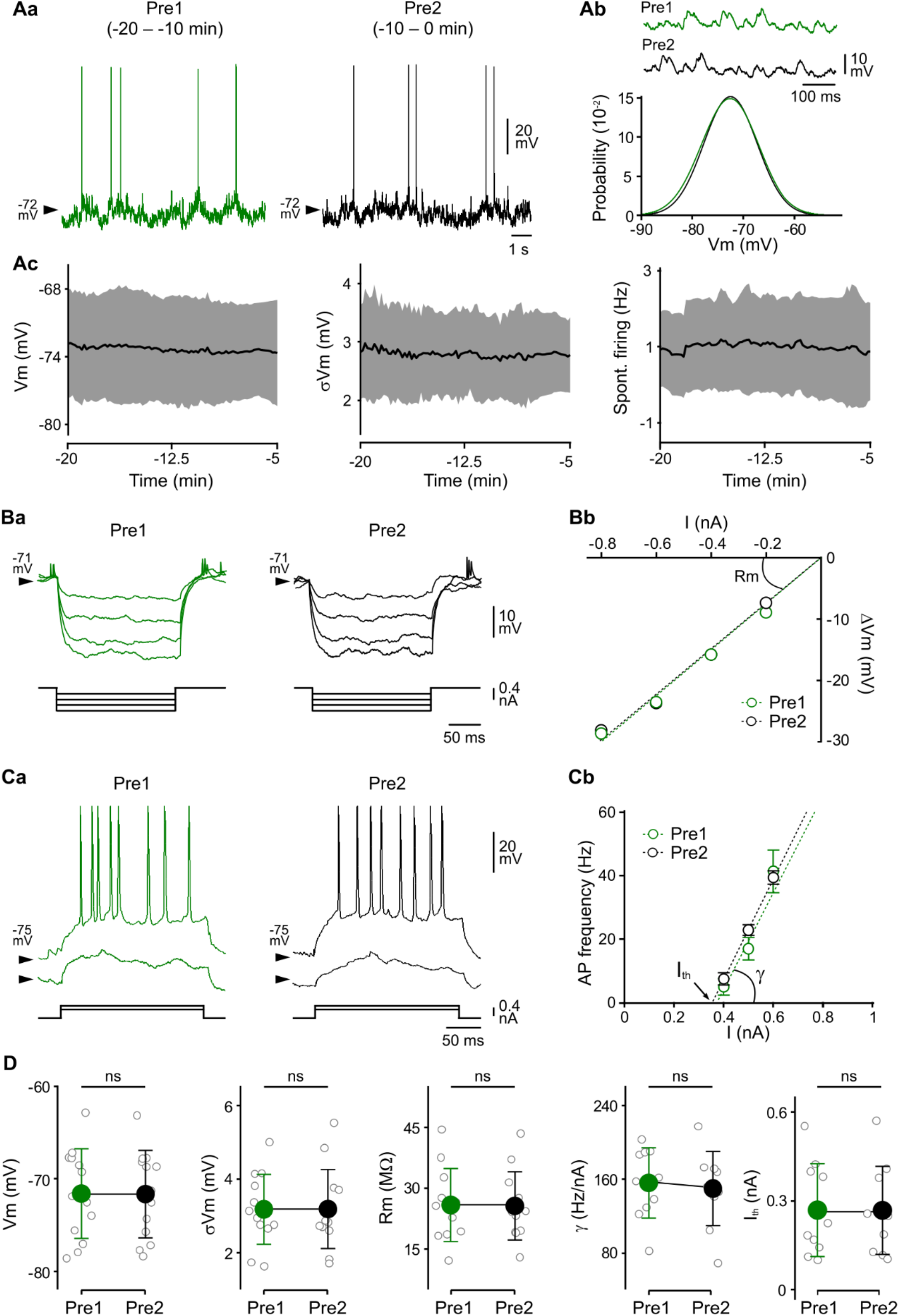
Stability of resting integrative properties and intrinsic excitability of cortical neurons over baseline periods. **(Aa)** Spontaneous intracellular activity of an S1 neuron recorded for two successive periods of 10 minutes: between 20–10 min (green trace, Pre1) and 10–0 min (black trace, Pre2) before the onset of the 10 min 10 Hz LI-rTMS protocol. Neuronal activity was characterized by a barrage of small-amplitude, broadband frequency, synaptic fluctuations (see also Fig. 2, A and B), confirming that sufentanil sedation induced cortical dynamics comparable to those observed during the waking state (Bruno and Sakmann, 2006; Constantinople and Bruno, 2011; Altwegg-Boussac et al., 2014). This background synaptic activity induced firing in 7 out of 13 neurons. Spontaneous firing rate of the active neurons was 2.9 ± 3.5 Hz during the first and 2.8 ± 3.5 Hz during the second pre-stimulation period (paired Student t-test, *P* = 0.29). Vm values are indicated at the left of intracellular records. **(Ab)** The probability densities of Vm values (60 s of recording, bin size 1 mV) demonstrate stability in the amplitude of synaptic activity over time. (Pre1 Vm = -71.6 ± 4.9 mV, *n* = 13 neurons *versus* Pre2 Vm, -71.7 ± 4.7 mV, *n* = 13 neurons; paired Student t-test, *P* = 0.92). **(Ac)** Time course of the mean values (solid lines, *n* = 8 neurons) ± SD (gray shaded areas) of Vm, σVm and spontaneous firing frequency computed over successive 30-second epochs during the 15 minutes of recording preceding LI-rTMS onset. Pearson’s correlations indicate a lack of significant changes over time. **(B)** Stability of membrane input resistance. **(Ba)** Average (*n* = 10– 17 trials) voltage responses (top traces) to negative current pulses of increasing intensity (bottom traces) recorded from an S1 pyramidal neuron during the two baseline periods (green traces, Pre1 and black traces, Pre2). The corresponding *V–I* relationships shown in **(Bb)** were best fitted by linear regressions (dashed lines, both *r²* > 0.99), indicating a lack of substantial membrane rectification in the hyperpolarizing direction. **(C)** Constancy of cell intrinsic excitability over the pre-stimulation period. **(Ca)** Examples of current-induced firing responses to depolarizing current steps of increasing intensity recorded during the two successive baseline periods (green traces, Pre1; black traces, Pre2). **(Cb)** *F-I* relationships and corresponding linear fits for the neuron illustrated in **(Ca)**. Each data point is the mean (±SD) firing rate calculated from 10–18 successive trials. **(D)** Population data comparing Vm, σVm, Rm, γ and I_th_ values during the two successive baseline periods. Gray circles correspond to values of individual neurons and green and black symbols show the mean ± SD. [Vm: *n* = 13; σVm: *n* = 13; Rm: *n* = 12; γ: *n* = 10; Ith: *n* = 10; paired Student t-test; \**P* < 0.05, *** *P* < 0.001, ns: non-significant].

**Fig. S2.**
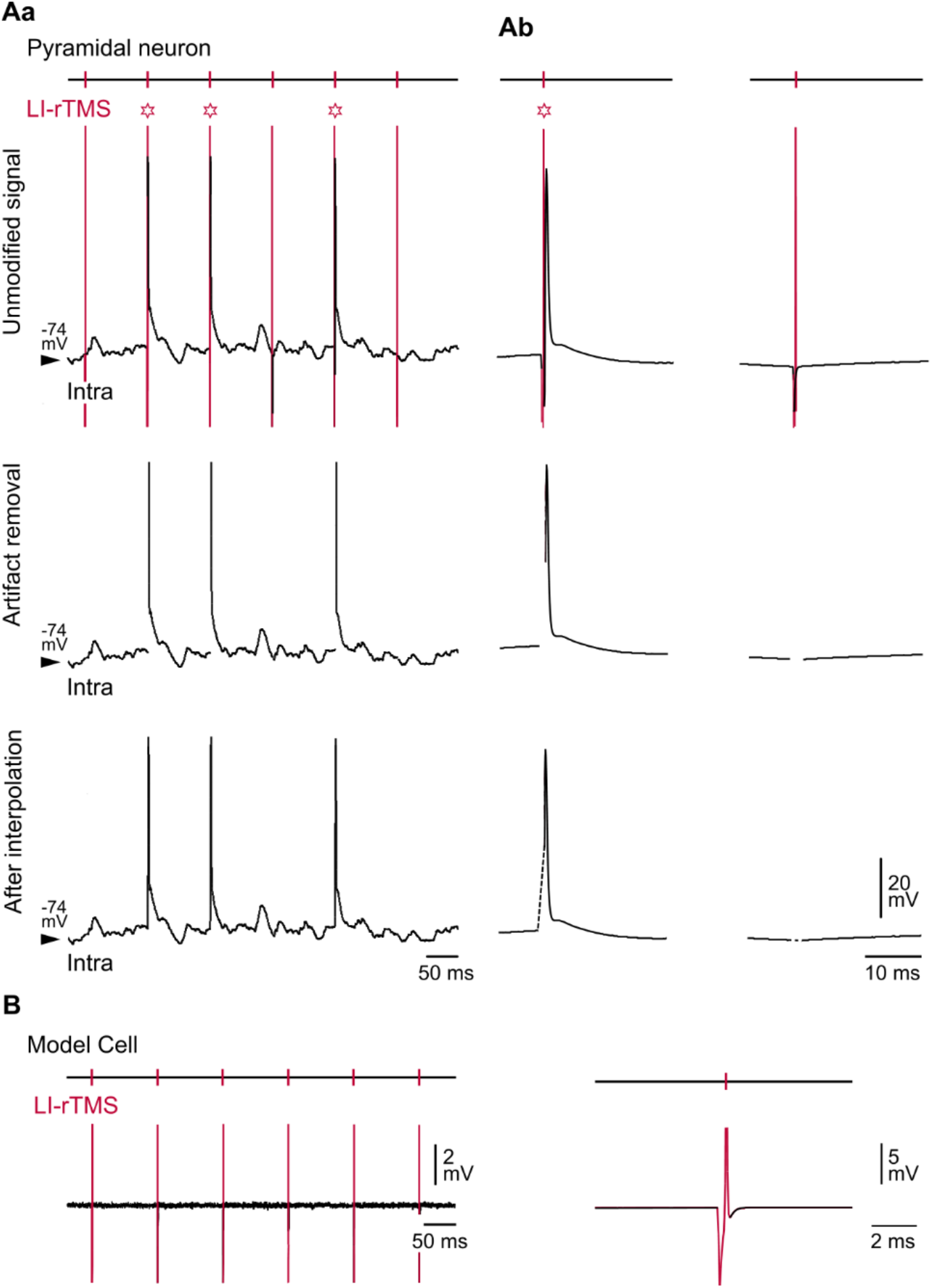
Procedure for removing LI-rTMS artifacts. **(A)** Off-line cutting and interpolation of electrical artifacts. **(Aa)** Short periods of intracellular activity recorded from a layer 2/3 pyramidal neuron during LI-rTMS, before removal of stimulation artifacts (vertical red lines, Unmodified signal), after their removal (Artifact removal), and after their replacement by linear interpolation (After interpolation). Red stars mark the onset of LI-rTMS-induced AP occurrences. Vm values are indicated at the left of the records. **(Ab)** Left, Representative example of an averaged (*n* = 2800) LI-rTMS-induced AP before and after removal of the LI-rTMS pulse stimulation artifact. Right, average cell responses (*n* = 3200) to magnetic pulses in absence of evoked APs. **(B)** Current-clamp recording during LI-rTMS illustrating the absence of evoked electrical responses in the aftermath of stimulation artifacts (in red) when a CLAMP-1U model cell is simulating the recording of a pyramidal neuron. An expanded view of a single stimulation is shown at right.

**Fig. S3.**
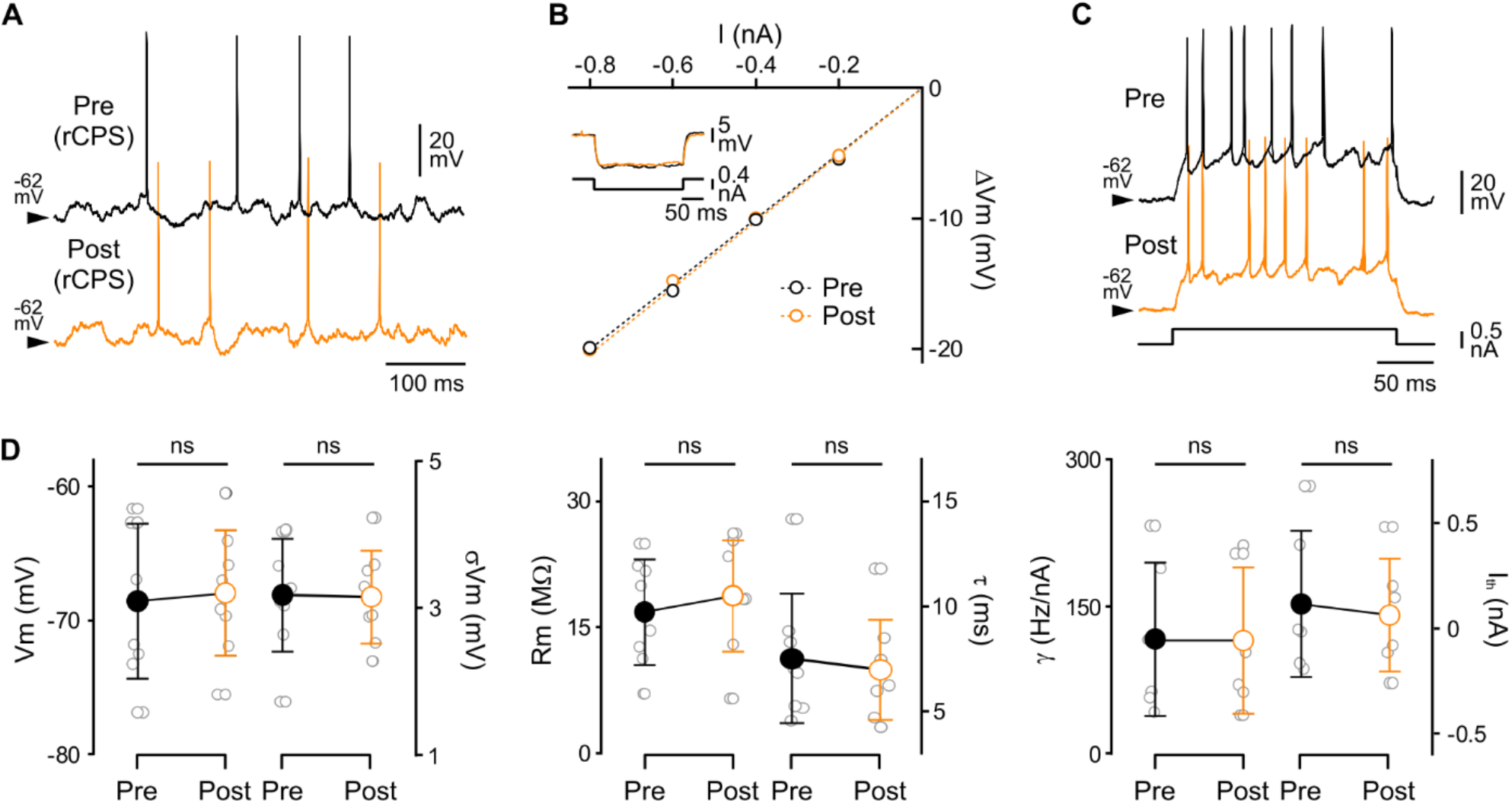
Stability of neuronal resting integrative properties and intrinsic excitability after rCPS. **(A**) Spontaneous intracellular activity recorded from an S1 neuron before (black trace, Pre) and 15 min after (orange trace, Post) the onset of the rCPS protocol. Vm values are indicated at the left of intracellular records. (**B**) *V–I* relationships computed from the neuron illustrated in (A) before and following the application of the rCPS protocol. The average voltage responses (20–30 trials) to negative current pulses of -0.4 nA (200 ms duration) during pre- and post-stimulation periods are superimposed in the inset. *V–I* relationships were best fitted by linear regressions (dashed colored lines, *r²* > 0.99). (**C**) Examples of current-evoked firing responses in baseline (Pre) and following rCPS (Post). (**D**) Population data comparing Vm, σVm, Rm, τm, γ and I_th_ values during the pre- and post-rCPS periods. Gray circles correspond to values of individual neurons and black and orange symbols show the mean ± SD. [Vm: *n* = 7; σVm: *n* = 7; Rm: *n* = 6; τm: *n* = 6; γ: *n* = 6; Ith: *n* = 6; paired Student t-test, ns: non-significant].

**Table S1:**
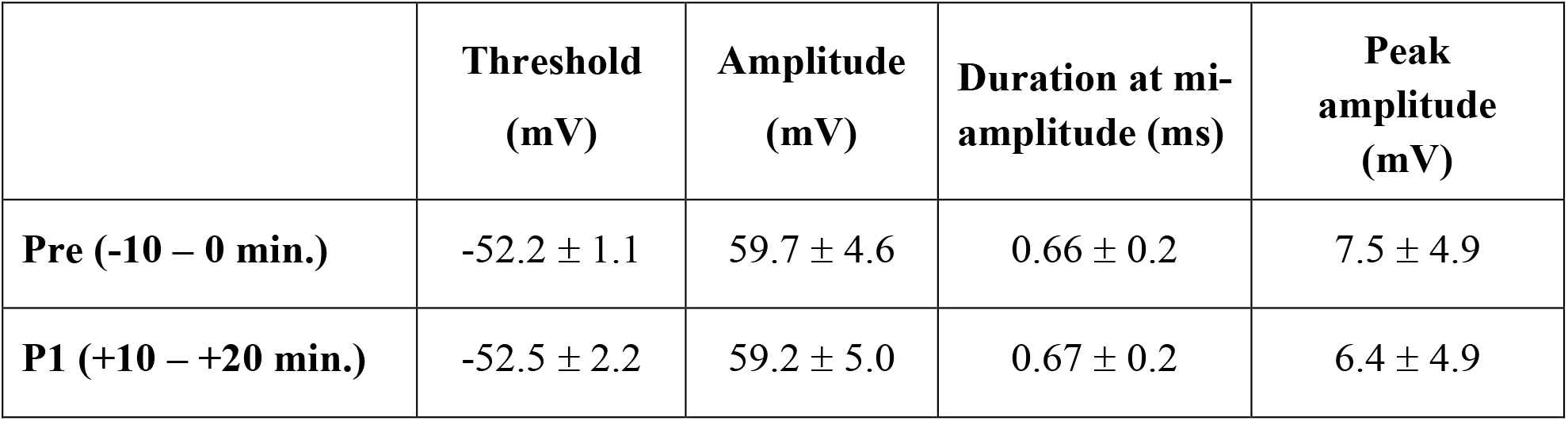
Properties of spontaneous APs are similar before and after LI-rTMS. Values are mean ± SD. No significant difference was found between the pre- and post-stimulation (P1) period (*n* = 8 neurons; paired Student t-test; *P* > 0.16 for each parameter).

