## Supplemental figures S1-S3 and Table S1 for "*In vivo* low-intensity magnetic pulses durably alter neocortical neuron excitability and spontaneous activity"

**This PDF file includes:**

Figs. S1 to S3
Table S1

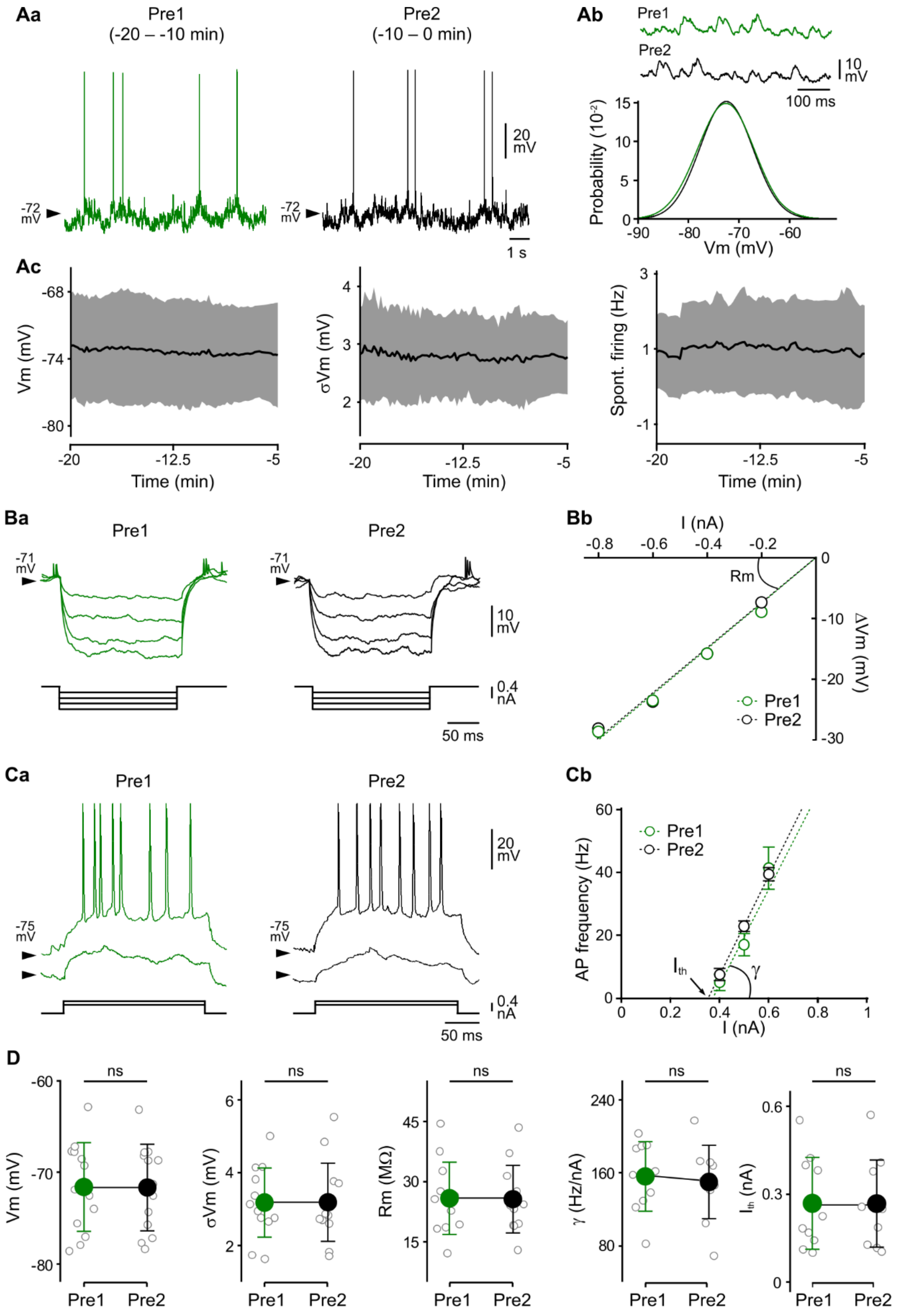
Aa
Pre1
(-20 – -10 min)
Pre2
(-10 – 0 min)
-72 mV
20 mV
1 s
Ab
Pre1
Pre2
10 mV
100 ms
Probability (10⁻²)
Vm (mV)
Ac
Vm (mV)
Time (min)
σVm (mV)
Spont. firing (Hz)
Ba
Pre1
Pre2
-71 mV
10 mV
0.4 nA
50 ms
Bb
I (nA)
Rm
ΔVm (mV)
Pre1
Pre2
Ca
Pre1
Pre2
-75 mV
20 mV
0.4 nA
50 ms
Cb
Pre1
Pre2
AP frequency (Hz)
I (nA)
Ith
γ
D
ns
Vm (mV)
σVm (mV)
Rm (MΩ)
γ (Hz/nA)
Ith (nA)
Pre1
Pre2

**Fig. S1. Stability of resting integrative properties and intrinsic excitability of cortical neurons over baseline periods.**

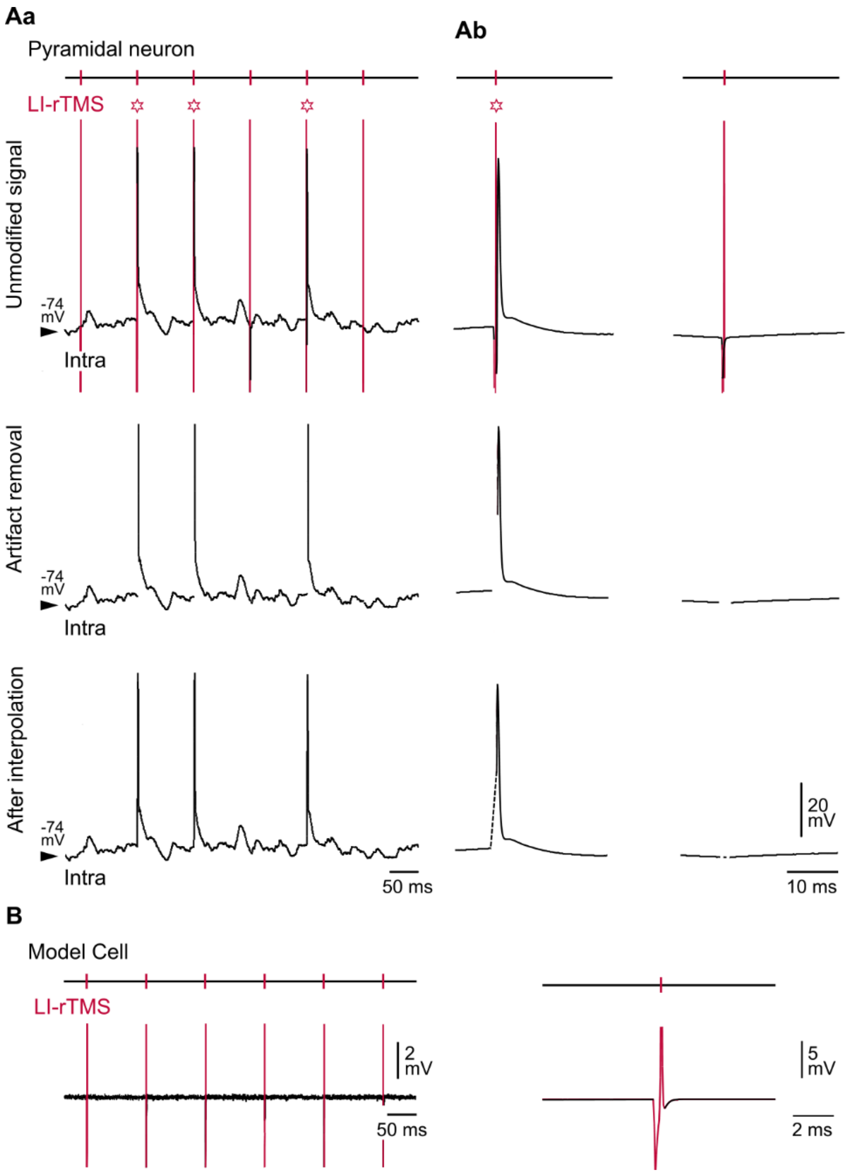


**Fig. S2. Procedure for removing LI-rTMS artifacts.**

**(A)** Off-line cutting and interpolation of electrical artifacts. **(Aa)** Short periods of intracellular activity recorded from a layer 2/3 pyramidal neuron during LI-rTMS, before removal of stimulation artifacts (vertical red lines, Unmodified signal), after their removal (Artifact removal), and after their replacement by linear interpolation (After interpolation). Red stars mark the onset of LI-rTMS-induced AP occurrences. Vm values are indicated at the left of the records. **(Ab)** Left, Representative example of an averaged ($n$ = 2800) LI-rTMS-induced AP before and after removal of the LI-rTMS pulse stimulation artifact. Right, average cell responses ($n$ = 3200) to magnetic pulses in absence of evoked APs. **(B)** Current-clamp recording during LI-rTMS illustrating the absence of evoked electrical responses in the aftermath of stimulation artifacts (in red) when a CLAMP-1U model cell is simulating the recording of a pyramidal neuron. An expanded view of a single stimulation is shown at right.

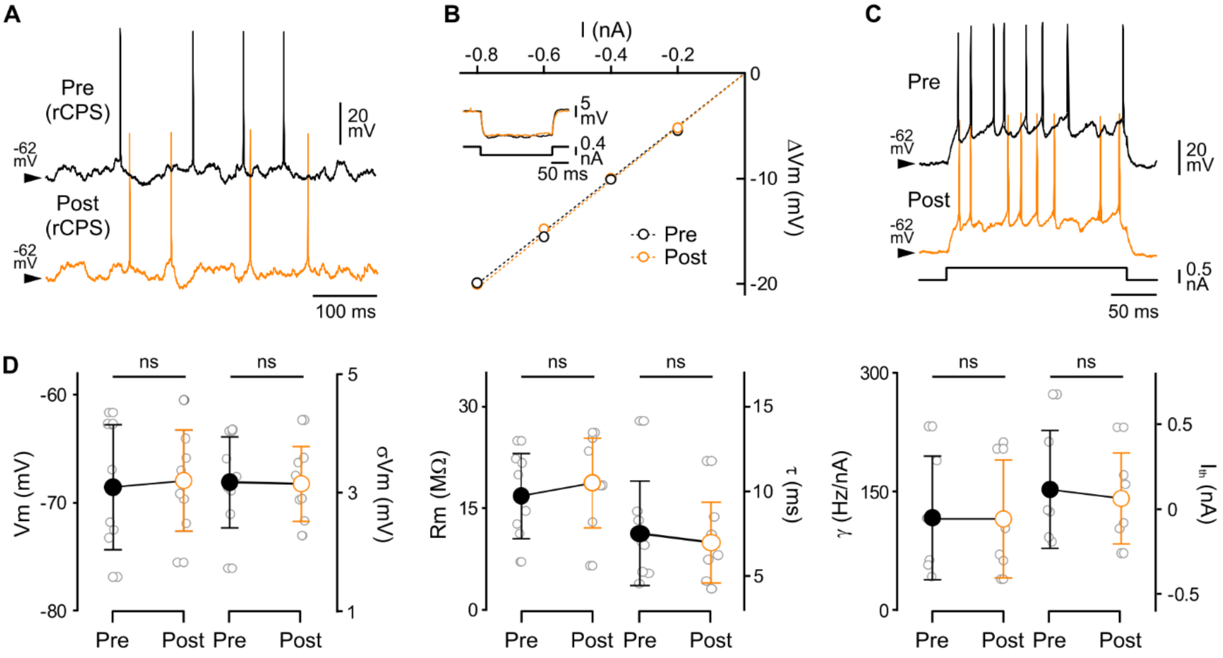


**Fig. S3. Stability of neuronal resting integrative properties and intrinsic excitability after rCPS.**

(**A**) Spontaneous intracellular activity recorded from an S1 neuron before (black trace, Pre) and 15 min after (orange trace, Post) the onset of the rCPS protocol. Vm values are indicated at the left of intracellular records. (**B**) *V–I* relationships computed from the neuron illustrated in (A) before and following the application of the rCPS protocol. The average voltage responses (20–30 trials) to negative current pulses of -0.4 nA (200 ms duration) during pre- and post-stimulation periods are superimposed in the inset. *V–I* relationships were best fitted by linear regressions (dashed colored lines, $r^2 > 0.99$). (**C**) Examples of current-evoked firing responses in baseline (Pre) and following rCPS (Post). (**D**) Population data comparing Vm, σVm, Rm, τm, γ and $I_{th}$ values during the pre- and post-rCPS periods. Gray circles correspond to values of individual neurons and black and orange symbols show the mean ± SD. [Vm: $n = 7$; σVm: $n = 7$; Rm: $n = 6$; τm: $n = 6$; γ: $n = 6$; Ith: $n = 6$; paired Student t-test, ns: non-significant].

| | Threshold (mV) | Amplitude (mV) | Duration at mi-amplitude (ms) | Peak amplitude (mV) |
|---|---|---|---|---|
| **Pre (-10 – 0 min.)** | -52.2 ± 1.1 | 59.7 ± 4.6 | 0.66 ± 0.2 | 7.5 ± 4.9 |
| **P1 (+10 – +20 min.)** | -52.5 ± 2.2 | 59.2 ± 5.0 | 0.67 ± 0.2 | 6.4 ± 4.9 |

**Table S1: Properties of spontaneous APs are similar before and after LI-rTMS.**

Values are mean ± SD. No significant difference was found between the pre- and post-stimulation (P1) period ($n$ = 8 neurons; paired Student t-test; $P > 0.16$ for each parameter).
